# One Circuit, Many Flow Fields: Mechanistic Models of Single-Trial Neural Dynamics

**DOI:** 10.64898/2026.06.01.729208

**Authors:** Shani Kaminitz, Mussi Levin, Ulises Pereira-Obilinovic, Ran Darshan

## Abstract

A single neural circuit can exhibit qualitatively different dynamics across trials: one circuit, many flow fields. Standard models treat this trial-to-trial variability as noise around a fixed dynamical system or as discrete switches between regimes, yet neither captures how continuous internal-state variables, such as arousal or engagement, can gradually deform the circuit’s flow field. We propose that single-trial fitting can be reframed as inferring the low-dimensional control parameters that reshape a shared circuit’s flow field. We realize this with a low-rank recurrent network in which trial-specific static input biases act as bifurcation parameters: constant within a trial, they deform the flow field without directly driving activity over time. We validate the model in a teacher-student setting, where it recovers the ground-truth dynamics and bifurcation structure from activity alone. Applied to mouse and monkey motor cortex, the model infers control spaces with interpretable axes that track behavioral engagement in mouse and task structure in monkey. A generative extension reproduces the structure of trial-to-trial variability and partially transfers across sessions and subjects. Together, these results reframe a methodological problem of fitting single-trial activity as a scientific opportunity: reading off the control parameters of the underlying dynamics, and connecting data-driven inference of neural dynamics to mechanistic theories of how a single circuit reuses its dynamics for flexible behavior.

## 1 Introduction

Neural circuits are often described as dynamical systems: population activity evolves through a flow field whose geometry reflects the computation being performed [Amari, 1977, Hopfield, 1982, Wong and Wang, 2006, Mante et al., 2013, Khona and Fiete, 2022]. In this view, fixed points, attractors, manifolds, and transient dynamics provide a compact language for explaining how circuits maintain memories, integrate evidence, or prepare movements. Yet this language is usually applied as if the relevant dynamical system were fixed across trials. Neural population activity, on the other hand, varies substantially from trial to trial [Engel et al., 2016, Shimaoka et al., 2018, Speed et al., 2020]. In the dynamical system perspective, this variability is then treated as variability *within* a flow field: different initial conditions, stochastic fluctuations, or noisy inputs drive different trajectories through the same underlying landscape [Pandarinath et al., 2018, Keshtkaran et al., 2022, Pals et al., 2024].

This picture is incomplete. In large-scale neural recordings, variability across trials is often structured, low-dimensional, and linked to slowly varying behavioral state [Steinmetz et al., 2019, Cowley et al., 2020, Wyrick and Mazzucato, 2021, Stringer et al., 2019]. Animals drift in engagement, arousal, movement state, expectation, or strategy, and these changes can alter how the same circuit responds to the same task events [McGinley et al., 2015, Ashwood et al., 2022, Allen et al., 2019, Fustiñana et al., 2021, Andalman et al., 2019]. From a dynamical-systems perspective, this suggests a stronger possibility: the circuit is not merely initialized differently on each trial, but may implement a different flow field. A single neural circuit may therefore generate a continuum of related dynamical systems—one circuit, many flow fields.

This distinction matters because different forms of variability imply different mechanisms. Noise around a fixed dynamical system changes where trajectories go, but not the rules that govern them. Switching models allow the rules to change, but typically between a small number of discrete linear regimes [Linderman et al., 2017]. Neither view captures how continuous, low-dimensional variables, such as arousal, engagement, or other modulatory signals, can gradually deform the vector field. The central question is therefore not only how to fit single-trial neural activity, but how to infer the low-dimensional parameters that reshape the circuit dynamics across trials.

Inferring trial-dependent flow fields from neural data is challenging because recordings provide sparse trajectories rather than direct observations of high-dimensional vector fields. This motivates a model that (i) preserves a shared recurrent scaffold across trials, (ii) captures trial-to-trial variability through low-dimensional continuous variables linked to internal or behavioral state, (iii) reveals how these variables reshape the dynamical landscape, and (iv) defines these variables as inputs to the recorded neurons, so that changes in them correspond to perturbations of the circuit that can be tested experimentally.

Our central contribution is a framework that goes beyond reconstructing single-trial activity with the goal of inferring a low-dimensional *control space*: the coordinates along which the flow field of a shared recurrent circuit is deformed across trials. We instantiate this framework with a low-rank recurrent neural network whose connectivity is fixed across trials, while a static, low-dimensional input bias selects one flow field from a continuous family generated by that connectivity (Fig. 1a). These biases can represent slowly varying long-range or neuromodulatory inputs and act as control parameters that shift fixed points, alter their stability or basin geometry, and induce bifurcations (Fig. 1b). Because each network unit corresponds to a recorded neuron and each bias is an input current onto those neurons, every direction in the inferred control space has two linked meanings: it describes how the latent flow field changes and specifies a concrete input pattern over identified cells. This dual role makes the model mechanistic rather than merely descriptive: it explains how trial-dependent inputs reshape recurrent dynamics and predicts which neurons should be driven, and how strongly, to move the circuit between dynamical regimes. The inferred mechanism can therefore be studied using low-rank network theory [Mastrogiuseppe and Ostojic, 2018] and tested in vivo using holographic photostimulations [Adesnik and Abdeladim, 2021, Finkelstein et al., 2026].

**Figure 1.**
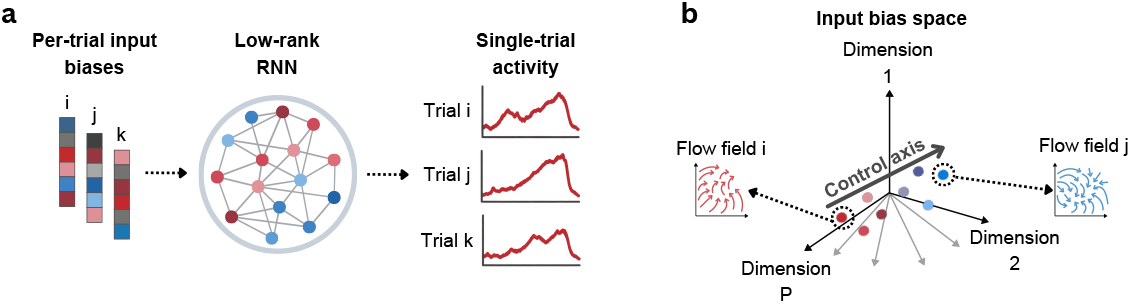
One circuit, many flow fields. (a) A low-rank RNN is fit to single-trial activity, with connectivity shared across trials and a static input bias inferred for each trial. (b) Each trial is a point in bias space that defines a flow field of the shared circuit; moving along a control axis continuously deforms the flow field.

### Contributions

- **A control-space model of trial variability**. We introduce a low-rank RNN in which continuous trial-specific inputs deform the flow field of a shared recurrent circuit, and infer this control space from single-trial population activity.
- **Ground-truth validation**. On synthetic data, the method recovers trial-dependent flow fields and their control space (bifurcation structure) from activity alone.
- **Internal-state control**. Applied to mouse motor cortex data, the model identifies a control axis for disengagement whose in-silico manipulation switches the fitted circuit between engaged and disengaged regimes.
- **Task-variable control**. We apply the framework to recordings from monkey motor cortex and identify a reach-direction control plane that generalizes to held-out directions; moving within this plane predictably rotates the decoded reach.
- **Shared low-rank dynamics across subjects**. We develop a generative extension of the model, which captures the distribution of single-trial activity, and its latent dynamics and variability structure transfer across subjects.

### Related Work

Our framework brings together two established lines of work: latent dynamical models that infer trial-specific inputs from single-trial activity, and analytically tractable low-rank recurrent networks whose units are the recorded neurons. Their combination lets trial-to-trial variability be read as the control parameters of a mechanistic circuit model.

#### Latent dynamical models

Sequential autoencoders such as LFADS [Pandarinath et al., 2018, Keshtkaran et al., 2022], neural ODE and state-space models [Kim et al., 2021, Schimel et al., 2021], Gaussian process approaches [Rutten et al., 2020], and deep recurrent models [Kim et al., 2023b, Luo et al., 2025] attribute trial-to-trial variability to initial conditions, inferred inputs, or stochastic fluctuations within latent dynamics shared across trials. Other latent models let the dynamics themselves change: switching models [Linderman et al., 2017] allow transitions among discrete regimes, and meta-dynamical state-space models map embeddings of individual recordings onto low-rank changes in the latent dynamics [Vermani et al., 2025]. Closest in form to our approach, Shah et al. [2025] provide the LFADS generator with an inferred per-trial bias, modeled as a constant input; the key difference is the system the bias enters, a generator of latent dynamics rather than a mechanistic circuit model of the recorded neurons.

#### Label-guided analyses

CEBRA [Schneider et al., 2023] learns embeddings of population activity that can be guided by behavioral variables, and MILCCI [Mudrik et al., 2026] captures cross-trial variability by decomposing multi-trial activity into components whose per-trial composition is adjusted according to trial labels. Both characterize how labeled variables are encoded in activity, without modeling the dynamics that generate it. Our per-trial inputs instead specify how a circuit’s dynamics are reshaped, and the behavioral variables we relate them to, such as engagement or reach direction, are not used as labels during training.

#### Low-rank circuit models

In low-rank recurrent networks, the dynamics reduce to a low-dimensional latent system whose fixed points, stability, and bifurcations can be analyzed directly [Mastrogiuseppe and Ostojic, 2018, Dubreuil et al., 2022]. Such networks have been inferred directly from neural recordings [Valente et al., 2022, Pals et al., 2024], but as a single dynamical system shared across trials. Pereira-Obilinovic et al. [2025] introduced the model we build on to study how activity along residual, non-task-coding dimensions drives choices; here we develop it into a general framework for inferring and interpreting the control parameters that reshape a circuit’s dynamics across trials.

#### Varying the circuit itself

Closest in spirit are models in which the recurrent weights themselves, and thus the circuit’s flow field, vary: Pellegrino et al. [2023] infer low-tensor-rank weight changes that track learning across trials, Costacurta et al. [2024] dynamically scale low-rank recurrent weights through a neuromodulatory subnetwork in task-trained networks, and JEDI [Jamkhandi et al., 2026] generates trial-specific recurrent weights from learned embeddings. In contrast to these methods, we ask how in a trained animal, a fixed circuit is reshaped from trial to trial by internal state or task demands, which calls for a different mechanism: low-dimensional changes in inputs rather than in synaptic weights. This also makes our model directly testable: inputs along the control axes can be delivered by targeted photostimulation [Adesnik and Abdeladim, 2021, Finkelstein et al., 2026], whereas prescribed changes to synaptic weights or gains have no experimental counterpart.

## 2 Model

We model single-trial neural population activity using a recurrent rate network with low-rank connectivity [Mastrogiuseppe and Ostojic, 2018, Pereira-Obilinovic et al., 2025], in which each recorded neuron corresponds to one neuron in the network [Rajan et al., 2016, Valente et al., 2022, Kim et al., 2023a, Pereira-Obilinovic et al., 2025]. Let *r*_*k*_(*t*) ∈ ℝ^*N*^ denote the firing rates of *N* neurons on trial *k* at time *t*. The network dynamics are given by

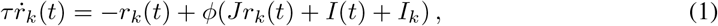

where *τ* is the neural time constant, *ϕ* is a bounded nonlinearity, 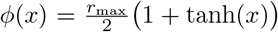, with learned per-neuron maxima *r*_max_ ∈ℝ^*N*^, *I*(*t*) denotes task-related inputs shared across trials, and *I*_*k*_ is a trial-specific static input.

### Low-rank recurrent connectivity

The connectivity *J*∈ ℝ^*N ×N*^ is constrained to be low-rank, *J* = *UAU*^*⊤*^, where *U*∈ ℝ^*N ×P*^ defines a *P* -dimensional latent subspace with *P*≪ *N*, and *A*∈ ℝ^*P ×P*^ specifies interactions between latent modes. Interactions between neurons are mediated through a low-dimensional latent space, while *A* governs the effective dynamics within it. The connectivity *J* is shared across all trials. Although neural activity evolves in the full *N* -dimensional space, the recurrent fields are confined to a low-dimensional subspace defined by *U* . This formulation supports a wide range of low-dimensional dynamics, including fixed-point attractors [Hopfield, 1982], sequential dynamics [Kleinfeld, 1986, Sompolinsky and Kanter, 1986], and continuous attractors [Seung, 1996, Darshan and Rivkind, 2022].

### Trial-specific input biases

Trial-to-trial variability is modeled through a low-dimensional bias vector *b*_*k*_ ∈ℝ^*P*^ for each trial, which enters the dynamics as a trial-specific static input *I*_*k*_ = *Ub*_*k*_. Crucially, *b*_*k*_ is constant within each trial and lies in the same low-dimensional subspace as the recurrent dynamics, ensuring that variability is parameterized along the same axes as the circuit’s intrinsic dynamics.

### Latent dynamics and bifurcation parameters

Projecting onto the latent subspace via *m*_*k*_(*t*) = *U*^*⊤*^*r*_*k*_(*t*) ∈ ℝ^*P*^, the dynamics can be written entirely in latent space:

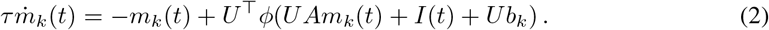

This defines a family of nonlinear dynamical systems on a shared *P* -dimensional latent space, parameterized by the trial-specific bias *b*_*k*_. Because *b*_*k*_ is constant within a trial, it acts not as a time-varying input but as a control parameter that deforms the flow field itself. Small changes in *b*_*k*_ smoothly reshape the dynamics, while critical values induce bifurcations that qualitatively reorganize the circuit’s response to the same task input *I*(*t*). Different trials therefore correspond to different dynamical landscapes generated by one common circuit (Fig. 1b).

### Fitting

We fit the model to recorded single-trial activity by jointly optimizing the parameters shared across trials, (*U, A, r*_max_) and, where present, those of the task input *I*(*t*), together with the per-trial biases{ *b*_*k*_*}* . The objective combines a dataset-specific data term with penalties that keep *U* approximately orthonormal and regularize the biases (Appendix A). To evaluate held-out trials, we freeze the shared parameters and optimize only the biases of those trials. See below for a generative model extension of this framework.

## 3 Results

### 3.1 Synthetic validation

We first validate the model in a controlled teacher-student setting, where the ground-truth dynamics and trial-specific inputs are known. Synthetic activity is generated from a rank-2 RNN generating bistable dynamics (Fig. 2a). In the absence of strong trial-specific inputs, the system exhibits two stable attractors corresponding to dominance of one of two neural populations. Each trial receives a constant low-dimensional bias input *b*_*k*_, which shifts the vector field and induces structured trial-to-trial variability: small changes to *b*_*k*_ modulate trajectories by shifting the boundaries between basins of attraction, while sufficiently large magnitudes eliminate one attractor, yielding a monostable regime (see Appendix C.1.1 for additional details).

**Figure 2.**
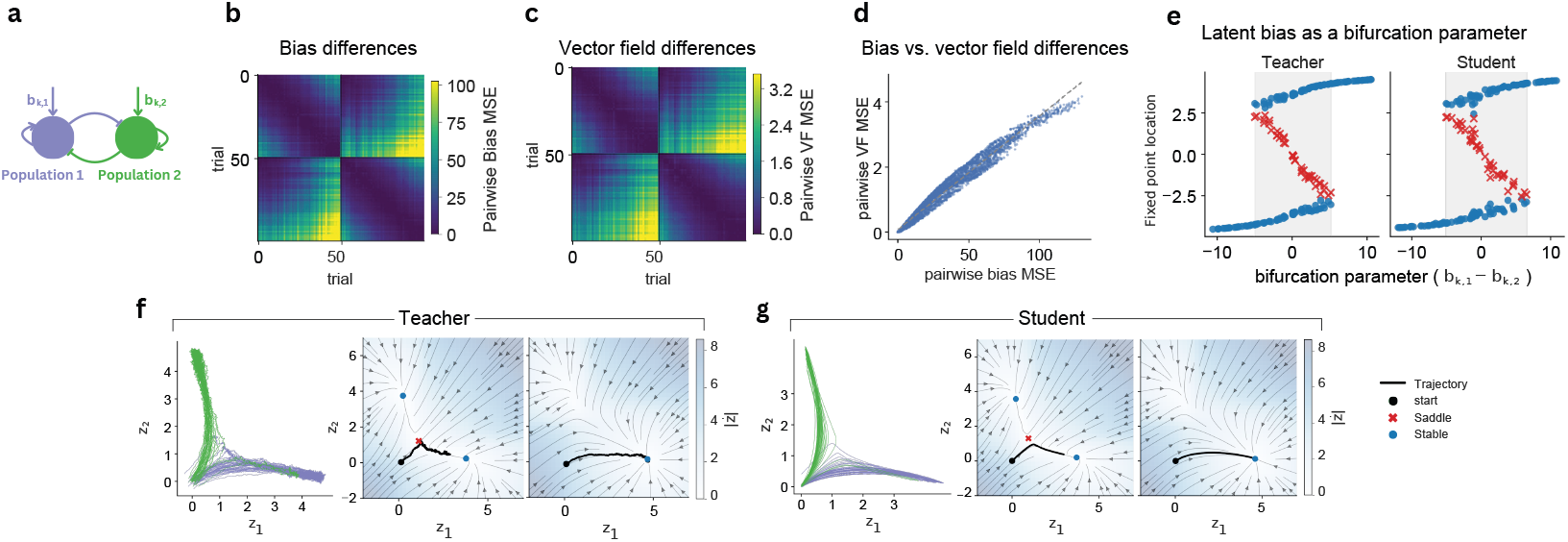
(a) Ground-truth system: a low-rank RNN (*P* = 2) with bistable dynamics and trial-specific bias inputs. (b,c) Pairwise distances between trials in bias space (b) and vector-field space (c) for the student model, computed as mean squared differences. (d) Pairwise differences in bias space versus the corresponding differences in vector-field space for the student model. (e) Fixed point locations as a function of the difference between the two latent bias dimensions. Shaded regions indicate the bistable regime, with two stable fixed points separated by a saddle. (f,g) Latent trajectories (left) and vector fields for representative trials (right), for the teacher (f) and student (g). For visualization, the latent spaces in (f,g) are aligned using a Procrustes transformation.

We then train a student model with the same architecture to reconstruct the teacher-generated activity. The trained model exhibits structured variability, as revealed by the pairwise differences between trials in both bias space and the induced vector fields (Fig. 2b,c). Moreover, variability in bias space is strongly correlated with variability in the corresponding vector fields (Fig. 2d), indicating that the model captures how low-dimensional shifts translate into changes in the dynamics. These pairwise distances also closely match those of the teacher (Fig. S1; Pearson *r* = 0.99).

Next, we examine how trial-specific biases reshape the fixed-point structure. In the teacher model, the bias difference *b*_*k*,1_ − *b*_*k*,2_ acts as a bifurcation parameter, inducing a transition from a bistable to a monostable regime (saddle-node bifurcation). The student reproduces this dependence, recovering both the number and locations of stable and unstable fixed points as a function of the bias difference (Fig. 2e). In addition, the student reconstructs both the neural and latent trajectories with high fidelity (Fig. 2; see also Appendix C.3 for details). Similar results hold for alternative ground-truth systems with different bifurcation structures (Fig. S3) and for students with overparameterized (larger *P* ) latent space (Fig. S2).

Together, these results show that the model recovers key features of the underlying dynamics and the structure of trial-specific variability directly from activity.

### 3.2 Reproducing single-trial activity in mouse motor cortex

We next test whether the model can reproduce single-trial neural activity in real cortical recordings. We apply it to population recordings from the anterior lateral motor cortex (ALM) of mice performing a delayed movement task [Chen et al., 2024], in which an auditory cue instructs a left or right lick after a delay period (Fig. S4).

To characterize task-related structure, we project activity onto coding dimensions, which provides a low-dimensional, interpretable summary of population dynamics. These are directions in population space that separate stimulus, choice, and response conditions [Chen et al., 2024, Pereira-Obilinovic et al., 2025] (Fig. 3a). For example, the choice dimension separates left and right trials during the delay period, reflecting the upcoming decision (Appendix D.2).

**Figure 3.**
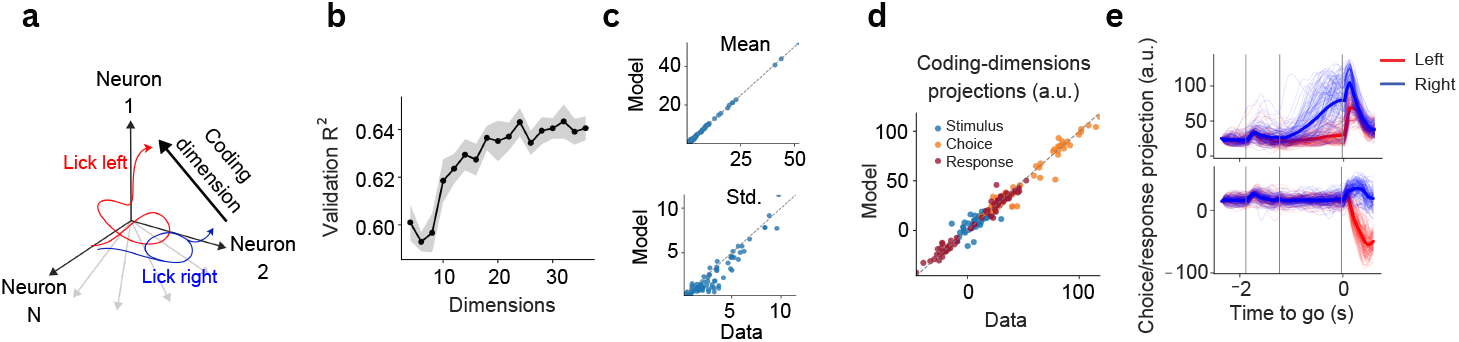
Reproducing single-trial activity in mouse ALM. (a) Schematic of coding dimensions. (b) Mean validation *R*^2^ vs. model dimensionality *P* (5-fold cross-validation on held-out trials. Error bars: standard error across folds). (c) Mean firing rates (in Hz; top) and standard deviations (Hz; bottom) across neurons in a session in data and model; dashed lines indicate identity. (d) Single-trial projections onto stimulus-, choice-, and response-coding dimensions in data and model, averaged over the corresponding task epoch; each point is one trial. (e) Single-trial activity (thin lines) projected onto choice (top) and response (bottom) coding dimensions for left (red) and right (blue) trials; thick lines: condition averages.

We apply the model to multiple recording sessions from the ALM dataset (Appendix D.1); results in this section are shown for a representative session. Validation performance increases with latent dimensionality and saturates at *R*^2^ ≈0.64 for *P* ≈20 (Fig. 3b), indicating that most predictive power is captured by a low-dimensional latent space. All results below are shown for *P* = 20; qualitatively similar results were obtained for nearby values of *P* .

On validation data, the model reproduces key population-level statistics (Fig. 3c,d). Quantitatively, the model achieves a global *R*^2^ of 0.64 on 5-fold validation data, with *R*^2^ values of 0.52, 0.83, and 0.73 for the stimulus, choice, and response projections, respectively. At the single-trial level, projections onto the choice and response dimensions show clear separation between behavioral conditions and the correct temporal evolution across task epochs (Fig. 3e). Although the model is not trained to fit individual neurons, it captures their single-trial responses (Appendix D.5); optimizing the full activity tensor instead improves neuron-level fits at the cost of accuracy along the coding dimensions (Appendix D.4). Together, these results show that modeling trial-to-trial variability as low-dimensional latent biases that modulate shared dynamics is sufficient to account for structured variability in cortical activity.

### 3.3 Control axis for engagement in mouse motor cortex

We next asked whether the inferred trial-specific biases can be used to identify control directions that reorganize the cortical dynamics across behavioral states. Although mice were highly trained, some sessions contained ignore trials in which the animal did not respond. For example, in the session shown in Fig. 4a, ignore trials became more frequent toward the end of the session, consistent with a gradual transition from an engaged to a disengaged behavioral state.

**Figure 4.**
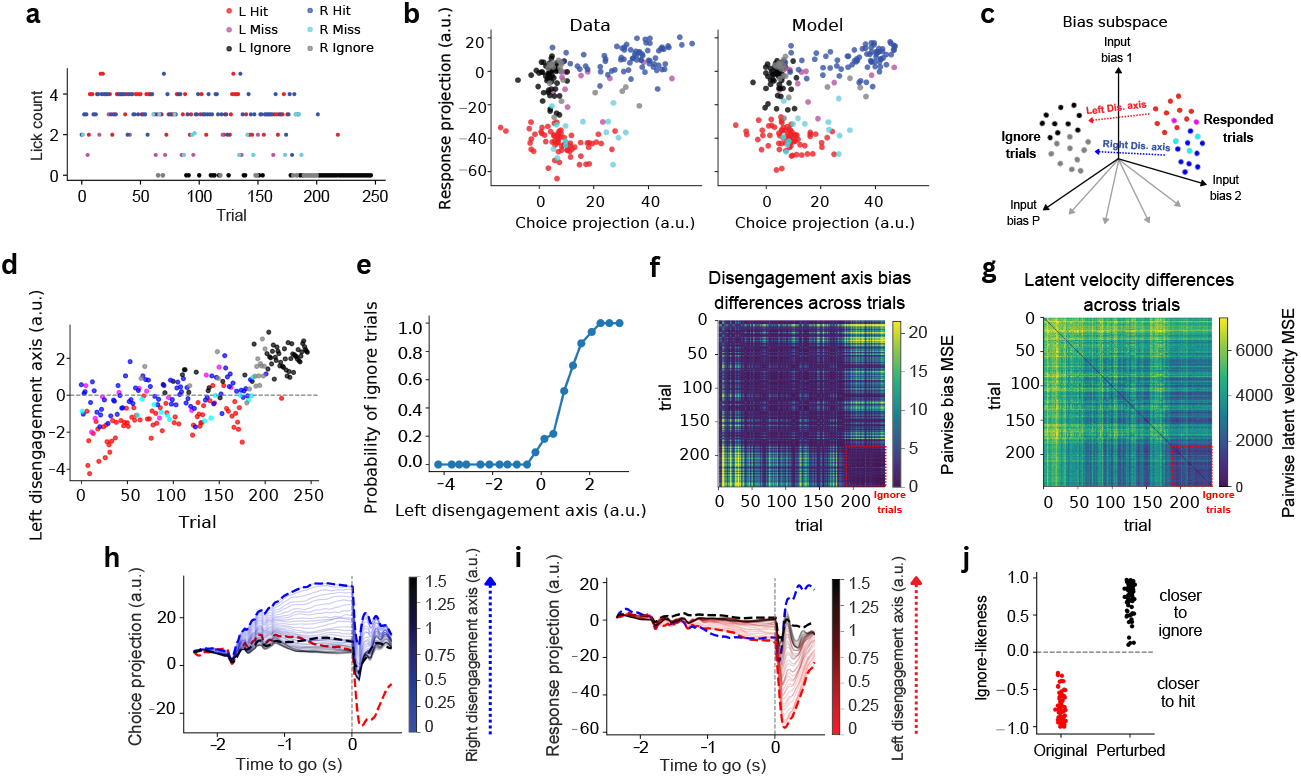
Control axis for disengagement. (a) Lick count across trials over the session. (b) Single-trial projections onto the choice and response coding dimensions, averaged over the delay epoch (choice) and response epoch (response). (c) Schematic of the disengagement axis in input-bias space (left and right). (d) Projection of per-trial biases onto the left disengagement axis over the session. (e) Probability of ignore trials vs. left-axis projection. (f) Pairwise differences between trials in bias space along the disengagement axis: squared difference of the trials’ projections onto the axis. (g) Differences between trials in latent velocity: squared difference between the time derivatives of their latent trajectories, averaged over time and latent dimensions. (h,i) Projections of neural activity onto choice (h) and response (i) following shifts along the right and left disengagement axes, respectively. (j) Single-trial ignore-likeness before and after a shift along the left disengagement axis (left hit trials). Ignore-likeness is defined as the normalized difference between distances to hit and ignore templates (dashed lines in h-i). Positive values indicate trajectories closer to the ignore template.

We trained the model on this session and analyzed the inferred trial-specific biases and latent dynamics. Ignore trials occupied distinct regions of neural state space: when projected onto the choice and response coding dimensions, ignore trials separated from responded trials in the data, and the model reproduced this separation (Fig. 4b). This separation suggests that behavioral disengagement is associated not merely with increased variability around a shared trajectory, but with systematic changes in the underlying dynamics.

To characterize these changes, for each stimulus condition (*c* ∈{ right, left} ) we defined a disengagement axis in bias space, by regressing the inferred biases on an ignore-versus-responded label (Appendix D.6). This yields an axis 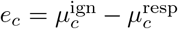 that estimates how the mean bias shifts from responded to ignore trials for the given condition. Importantly, these axes emerged entirely from the inferred control-parameter space of the model, behavioral labels were used only after model fitting.

Projecting all trials onto the disengagement axis revealed a continuous organization of behavioral state in the inferred bias space (Fig. 4d,e). Ignore trials occupied the extreme end of the axis, while responded trials formed a continuum extending toward them. In this session, disengagement-axis projections also increased gradually over the course of the experiment (Fig. 4d), consistent with a slow drift of the circuit toward a disengaged dynamical regime. The disengagement axes for left and right trials were partially aligned (cosine similarity = 0.53), suggesting that engagement modulates shared components of the dynamics while preserving condition-specific structure.

We next asked whether movement along the disengagement axis systematically reorganized the latent dynamics. As in the synthetic teacher-student system, nearby trials in disengagement-bias space induced similar latent dynamics. Pairwise differences between trials projected onto the disengagement axis revealed a distinct cluster of ignore trials (Fig. 4f). The same trials also exhibited similar latent velocity trajectories (Fig. 4g), indicating that nearby points in disengagement-bias space correspond to nearby dynamical regimes. In contrast, responded trials exhibited substantially greater variability in their latent velocity trajectories, reflecting a broader diversity of dynamics during engaged behavior. Thus, as in the synthetic examples, low-dimensional trial-specific biases organize a continuous family of related flow fields.

We then directly tested the dynamical role of the disengagement axis by shifting the inferred biases of responded trials along it, *b*_*k*_(*λ*) = *b*_*k*_ +*λe*_*c*_, and simulating the resulting activity with the recurrent connectivity fixed (Appendix D.6). Shifts along the disengagement axis systematically deformed the latent dynamics toward the ignore-trial regime. For right trials, shifts along the right disengagement axis progressively reduced projections onto the choice dimension (Fig. 4h), while for left trials, shifts along the left axis shifted trajectories along the response dimension toward ignore-like dynamics (Fig. 4i). Shifting ignore trials in the opposite direction (*λ <* 0) produced complementary effects, moving their trajectories toward the responded regime.

To quantify the effects of these shifts at the single-trial level, we defined an ignore-likeness metric as the normalized difference between the distance of a trajectory to responded and ignore templates. Shifts along the disengagement axis systematically increased ignore-likeness for responded trials (Fig. 4j), indicating that these continuous movements along the inferred control axis reshape the dynamics toward disengaged trajectories.

Together, these findings suggest that structured trial-to-trial variability reflects movement through a low-dimensional control space that deforms neural dynamics.

### 3.4 Generative modeling of trial variability reveals shared low-rank dynamics across mice

A central idea of our framework is that trial-to-trial variability reflects movement through a low-dimensional space of control parameters that deform a shared recurrent flow field. We therefore extended the model to a generative setting in which trial-specific biases are treated as random variables, with *b*_*k*_ ∼ *N*(0, Σ) (see full description of the generative model in Appendix D.3).

We trained a separate generative model for each session, using a trial-matching objective [Sourmpis et al., 2023] to optimize (*U, A*, Σ) so that generated activity reproduced the statistical structure of the recorded activity (Appendix D.3; Fig. 5b). Sampling latent biases therefore amounts to sampling from a distribution of flow fields generated by a single circuit, with Σ defining the geometry of trial-to-trial variability in latent control space.

**Figure 5.**
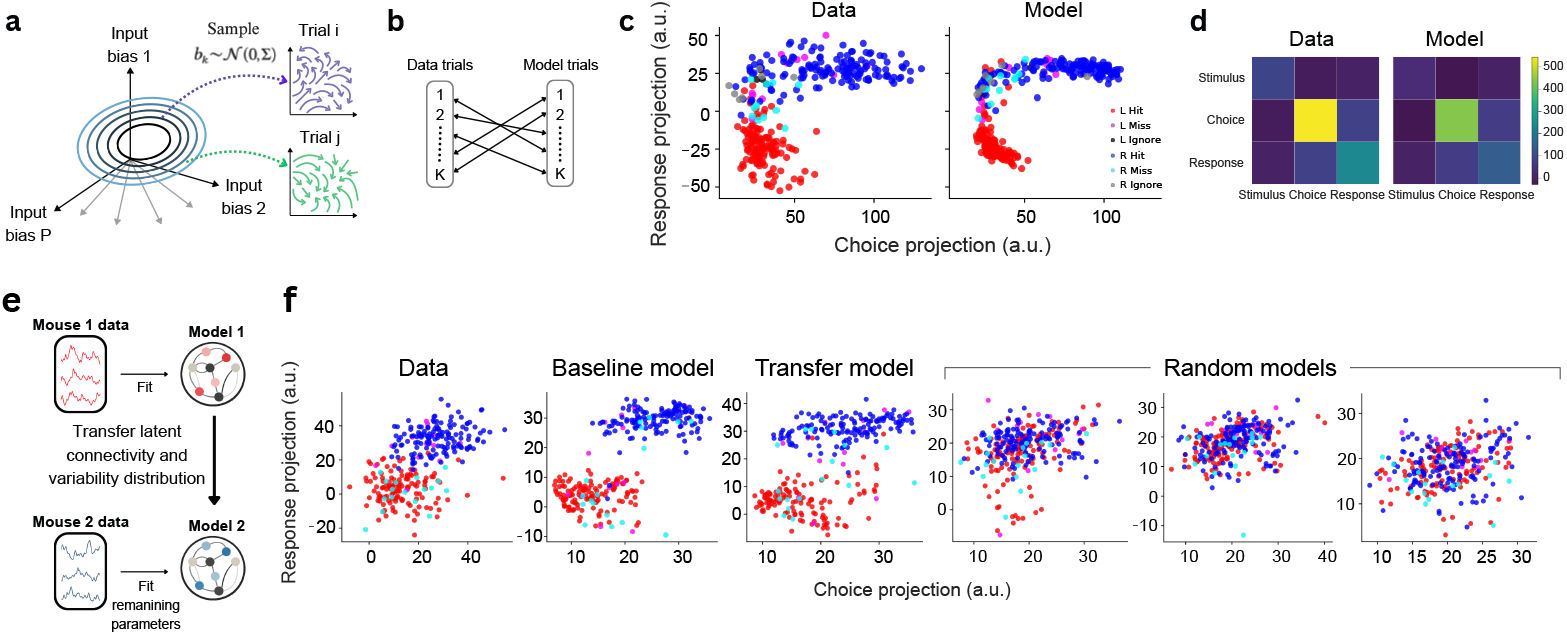
Generative model evaluation on held-out trials. (a) Generative model schematic. Per-trial biases are sampled from a learned distribution. (b) Trial-matching schematic, used to compare generated and recorded trials. (c) Single-trial projections onto the choice and response coding dimensions, averaged over the delay and response epochs respectively, for data and model-generated trials. (d) Covariance of coding-dimension projections for data and model. (e) Cross-session transfer schematic. Latent dynamics *A* and variability Σ learned from one session are transferred to another session while retraining the remaining parameters. (f) Single-trial projections onto choice and response coding dimensions for data, baseline, transfer, and random models.

The generative model reproduced the structure of single-trial variability across multiple measures: generated trials matched the distributions of single-trial activity statistics (Fig. S7), their covariance in coding-dimension space, and the clustering of trial outcomes (hit, miss, ignore), which were not used in training (Fig. 5c,d).

We next asked whether the learned latent dynamics generalize across recording sessions. To test this, we performed a cross-session transfer analysis between sessions recorded from different mice performing the same task (Fig. 5e; session IDs in Appendix D.1). We compared: (i) baseline models trained directly on the target session; (ii) transfer models in which the latent dynamics *A* and variability distribution Σ learned from a different session were kept fixed while retraining the remaining parameters; and (iii) random control models in which *A* and Σ were replaced with norm-matched random matrices.

Transfer models achieved validation performance close to baseline models and substantially better than random controls (Fig. S8). Moreover, transfer models reproduced the task-aligned clustering observed in coding-dimension space, whereas random models failed to recover this structure (Fig. 5f). Together, these results suggest that across animals performing the same task, trial-to-trial variability is organized by a shared low-dimensional control space that generates a continuum of related flow fields within one recurrent circuit.

### 3.5 Control plane for reach direction in monkey motor cortex

We next applied the framework to macaque motor and premotor cortex recordings during delayed reaches across 108 target and barrier configurations [Pei et al., 2021, Churchland et al., 2010]. Without task inputs, reach direction could enter only through the inferred biases, allowing us to test whether they recover interpretable task-related rather than internal-state control directions. On the Neural Latents Benchmark, our model outperformed classical latent-variable baselines on every reported metric and was close to substantially more expressive deep models (Appendix E.3).

Focusing on barrier-free trials, we asked whether reach direction is encoded in an interpretable, low-dimensional subspace of the bias space, and whether moving within that subspace steers the reach. To identify how reach direction is encoded in bias space, we regressed the inferred biases on target angle, yielding a two-dimensional reach-direction plane (Appendix E.4). To test generalization, we inferred this plane while excluding trials with target angles within ± 20° of 90° (*n* = 89). When projected onto the inferred plane, the biases formed a ring ordered by target angle, with the held-out trials falling at the appropriate location (Fig. 6a, outlined). As in the synthetic and ALM settings, nearby trials in this plane exhibited similar latent dynamics: pairwise distances in the reach-direction plane (Fig. 6c) were correlated with pairwise differences in latent velocity (Fig. 6d; *r* = 0.75).

**Figure 6.**
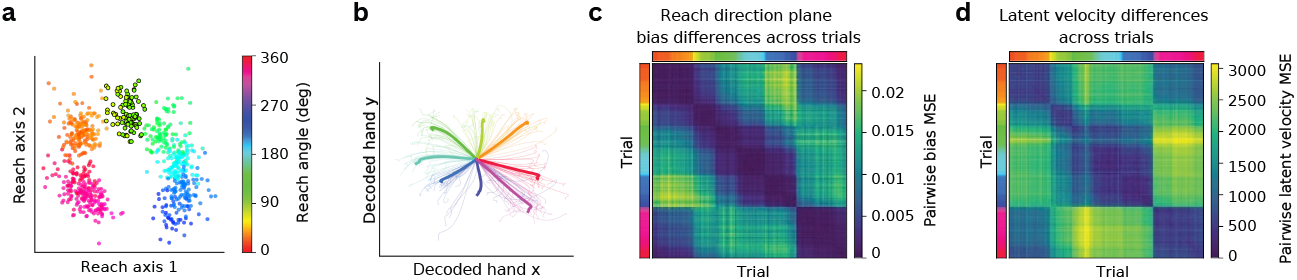
Reach-direction control plane in monkey motor cortex. (a) Inferred biases projected onto the reach-direction plane fit without trials within ±20° of 90° (outlined), colored by reach angle. (b) Decoded hand paths after shifting biases within the plane, for trials with targets within ±10° of 0° (*n* = 23). Colors indicate the imposed target angle; the original reach is red. Thick lines: mean over trials; thin lines: single trials. (c,d) Pairwise differences between trials in the reach-direction plane (c; mean squared difference of the trials’ plane coordinates) and in latent velocity (d; as in Fig. 4g). Trials are sorted by reach angle, shown by the color strips (colors as in a).

We then tested whether moving within the plane controls reach direction. For each barrier-free trial, we moved the bias within the plane from the position of its target angle, keeping the remaining components of the bias and the recurrent connectivity fixed, and decoded hand velocity from the simulated activity (Appendix E.4). Shifting the biases rotated the decoded reach by approximately the imposed angle (Fig. 6b).

Thus, without being given the task, the model organized reach direction as a continuous plane in its control space: nearby positions on the plane produce similar dynamics, and moving a trial along it steers the decoded reach. We note that also previous models [Pals et al., 2024, Shah et al., 2025] also recovered reach geometry on the same dataset, but Pals et al. supplied target position as an input and the biases of Shah et al. acted on an LFADS latent generator. In contrast, here, the reach-direction plane emerges without task input and maps directly to the dynamical system of the recorded neurons through *I*_*k*_ = *Ub*_*k*_. Like in the ALM dataset, this mapping allows a concrete prediction for how to experimentally steer the circuit, using patterned optogenetic stimulation [Finkelstein et al., 2026].

## 4 Discussion and limitations

Our framework recasts non-stationary trial-to-trial variability as movement along a small set of control axes that qualitatively reshape the circuit’s dynamics, rather than as independent noise to be averaged away. A central feature of the framework is that this variability is constrained to the same low-dimensional subspace as the recurrent dynamics. Two findings support this choice. First, when fit to mouse and monkey motor cortex, the framework recovers behaviorally meaningful axes. These axes are interpretable through their alignment with the recurrent subspace and causally shape the model’s dynamics. Trial-to-trial variability can therefore be understood as continuous deformations within the subspace where the recurrent dynamics live (see also Pereira-Obilinovic et al. [2025]). Second, connectivity and bias distributions inferred from one animal partially transfer to others, indicating that the subspace reflects shared structure across animals rather than quirks of a single recording. Together, these results suggest that control axes in input-bias space provide a coordinate system over a family of dynamics: moving along them deforms the circuit’s flow field and can drive bifurcations, which produce abrupt transitions in the neural dynamics underlying behavior.

### Limitations

Our framework treats trial-specific biases as independent across trials. Yet, in many paradigms (e.g. Bari et al. [2019], Vertechi et al. [2020], International Brain Laboratory et al. [2021], Rajagopalan et al. [2023]), biases on one trial might depend nonlinearly on the history of preceding trials. Capturing this requires additional dynamics for the biases themselves, which we leave to future work.

## AI Use Statement

Generative AI tools were used to help write parts of the code, which the authors reviewed and tested, and for writing assistance: editing the manuscript for readability and length, drafting appendix descriptions and captions, and checking wording and references. The authors reviewed all AI-assisted content and take full responsibility for the paper.

## Acknowledgments and Disclosure of Funding

We thank Kabir Dabholkar for helpful comments on the manuscript. This research was supported by the Israel Science Foundation (ISF, 2097/24 and 2994/24) and the TAU Center for Artificial Intelligence & Data Science (TAD). RD was supported by the Alon scholarship for the integration of outstanding faculty. UP was supported by the Allen Institute.

## A. Training setup

All models share the same architecture, regularization and optimization, differing only in the data term of the objective and a few per-dataset settings.

### Training objective

The training objective has the form

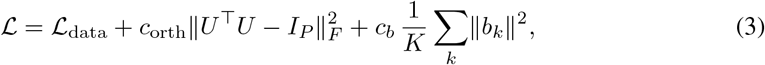

whereℒ _data_ is defined separately for each dataset (Appendices C.2, D.3, E.2). The orthogonality penalty keeps *U* approximately orthonormal, so that *m*_*k*_ are the coordinates of activity projected onto the recurrent subspace; the penalty on the biases regularizes the trial-specific parameters, limiting overfitting to trial-specific noise.

### Inference on held-out trials

Because each trial carries its own bias vector, evaluating unseen trials requires inferring new biases: we freeze the trained network parameters and optimize only the held-out biases {*b*_*k*_*}* under the same objective, so no information from held-out trials enters (*U, A, r*_max_).

### Optimization

All parameters were optimized with Adam (learning rate 5 ×10^*−*4^, gradient clipping at global norm 10), implemented in PyTorch 2.6 and trained on a single NVIDIA L40S GPU. Training a single model takes around 1 hour for the synthetic data, 5–6 hours for the ALM data, and 9 hours for the MC Maze data.

## B Encoding axes in bias space

To relate the inferred biases to task or behavioral variables, we regressed the biases on the variables. For trial *k* with variables *y*_*k*_ ∈ ℝ^*m*^, the regression is

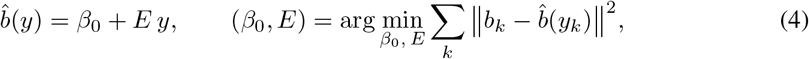

with intercept *β*_0_ ∈ ℝ^*P*^ and coefficients 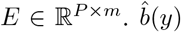 is the part of the bias explained by the variables, and the *j*-th column of *E*, the encoding axis of variable *j*, is the change in bias per unit change of *y*_*j*_. For a single binary variable coded as 0*/*1, the encoding axis is the difference between the mean biases of the two classes.

To perturb trial *k* as if its variables had taken the value *y*^*′*^, we replaced the explained part of its bias and left the rest unchanged:

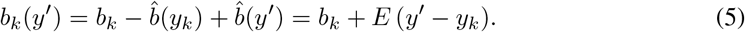

## C Synthetic networks

### C.1 Data generation

#### C.1.1 Bistable system

Synthetic data were generated from a rank-*P* =2 network with bistable dynamics (Table 1). The embedding matrix *U* ∈ ℝ^*N ×*2^ had a block structure defining two disjoint neural populations, with non-zero entries drawn from a log-normal distribution, normalized to unit norm, and shared across populations up to a random permutation. The interaction matrix *A* ∈ ℝ^2*×*2^ was constructed as:

**Table 1.** Hyperparameters for the synthetic networks.

|  | Bistable | Limit-cycle |
| --- | --- | --- |
| *Architecture* |  |  |
| Neurons $N$ | 100 | 100 |
| Latent dimensionality $P$ | 2 | 2 |
| Time constant $\tau$ | 1 s | 10 ms |
| Integration step $\Delta t$ | 10 ms | 1 ms |
| *Data generation* |  |  |
| Trials $K$ | 100 | 200 |
| Trial duration | 10 s | 1 s |
| Noise $\sigma$ | 0.1 | 0.2 |
| Firing-rate maximum $r_{\max}$ | 1.0 | 1.0 |

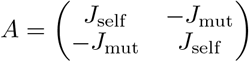

where *J*_self_ (*J*_self_ = 1.0) defines the self-excitation within each population, and *J*_mut_ (*J*_mut_ = 5.0) defines the mutual cross-inhibition between them. Trials were divided into two blocks each favoring a different population, with trial-specific biases held constant within each trial. Within each block, the bias for the favored population increased linearly from 1.0 to 11.5 across trials, while the competing population received 0.5 + *u*_*k*_ with *u*_*k*_ ∼ *U* (0, 1). Gaussian white noise was added at each time step to model unstructured variability.

#### C.1.2 Limit-cycle system

To test recovery under a different bifurcation structure, we generated a second ground-truth system: a rank-*P* =2 network with rotational latent structure and stable limit-cycle dynamics (Table 1). The columns of *U* ∈ ℝ^*N ×*2^ corresponded to the cosine and sine of evenly spaced angles in [0, 2*π*]. To generate these stable rotational dynamics, the interaction matrix *A* was constructed as:

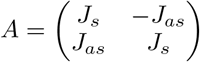

where the parameter *J*_*s*_ (*J*_*s*_ = 6.0) defines the symmetric connectivity that stabilizes the amplitude of the localized activity bump, and *J*_*as*_ (*J*_*as*_ = 3.0) determines the anti-symmetric connectivity that drives its continuous rotation. In the absence of trial-specific biases, the interaction matrix *A* was chosen to generate stable rotational dynamics in the latent plane.

Trials were divided into two blocks sampling distinct dynamical regimes. In block 1, both modes received biases drawn from *U* (0, 10), keeping the network in its limit-cycle regime. In block 2, the first mode was strongly driven with *b*_*k*,1_ *∼ U* (15, 25) while the second received *b*_*k*,2_ *∼ U* (0, 1), pushing the system into a fixed-point regime. Gaussian white noise was added at each time step to model unstructured variability.

### C.2 Training

The synthetic networks receive no task input (*I*(*t*) = 0). With 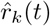denoting model-simulated rates, the data term in Eq. 3 is the mean squared error over the full activity tensor,

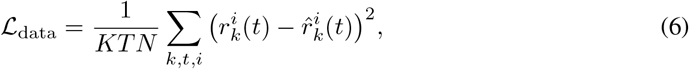

since the synthetic data are firing rates rather than spike counts and carry no task-defined coding dimensions. We used *c*_orth_ = 500 and *c*_*b*_ = 0.

### C.3 Latent space alignment

The low-rank parameterization *J* = *UAU*^*⊤*^ is invariant under orthogonal transformations of the latent space. Specifically, for any orthogonal matrix *Q*, the transformation *U* →*UQ* and *A* →*Q*^*⊤*^*AQ* leaves the full connectivity matrix *J* unchanged. As a result, when provided with only the neural activity, the latent representation is only identifiable up to rotation.

To compare student and teacher latent trajectories in Section 3.1, we align the student latent space to the teacher using an orthogonal Procrustes transformation. This amounts to finding the orthogonal matrix *Q*^*\**^ that minimizes ∥*U*_teacher_ −*U*_student_*Q* ∥_*F*_ . All comparisons of latent trajectories and parameters are performed after this alignment.

### C.4 Additional results for the bistable system

Pairwise trial distances in both bias and vector-field space are highly correlated between teacher and student (*r* = 0.99; Fig. S1), indicating that the student recovers the geometry of trial-to-trial variability.

We also examined the case in which the student’s latent space is overparameterized (*P >* 2). The model recovers the underlying two-dimensional dynamics, bifurcation structure and attractor geometry, with the first two principal components capturing over 99% of latent variance and additional dimensions contributing negligibly (Fig. S2).

### C.5 Recovery of limit-cycle dynamics

We repeated the teacher-student analysis of Section 3.1 on the limit-cycle system. The student model recovered the global bifurcation structure, properly transitioning from limit cycles to fixed-point dynamics (Fig. S3d). While the exact flow field is not always captured, variability in bias space reliably predicts variability in the corresponding vector fields (Fig. S3c). Pairwise trial distances in both bias and vector-field spaces are highly correlated between the teacher and student (*r* = 0.95, Fig. S3h, i), indicating that the model captures the geometry of the underlying dynamics.

## D Mouse ALM

### D.1 Data

All mouse analyses use sessions from the publicly available ALM dataset of Chen et al. [2024], where mice performed a delayed-response licking task (Fig. S4). During the sample epoch, one of two auditory tones instructed the direction of the upcoming lick. After a delay epoch, a go cue initiated the response epoch, in which the mouse reported its choice by licking left or right. Trials were classified as hit (correct lick), miss (lick in the wrong direction), or ignore (no lick after the go cue). The sample, delay, and response epochs define the coding dimensions (Appendix D.2) and the epoch averages in the training objective (Appendix D.3).

Spiking activity was binned at 10 ms and smoothed with a causal exponential kernel (50 ms time constant) to obtain firing rates. Tables 2 and 3 list the sessions used for each analysis.

**Table 2.**
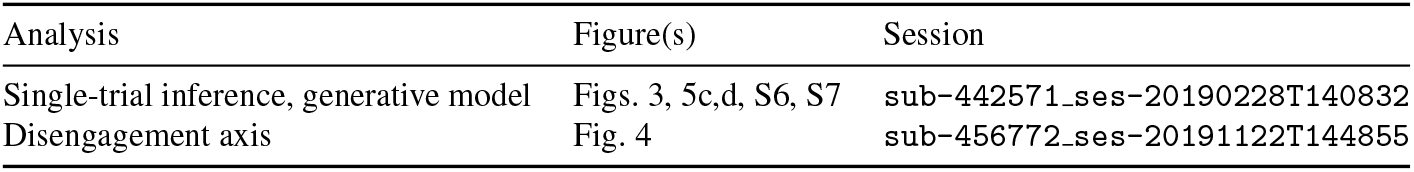
Sessions used for single-session analyses.

| Analysis | Figure(s) | Session |
| --- | --- | --- |
| Single-trial inference, generative model | Figs. 3, 5c,d, S6, S7 | sub-442571_ses-20190228T140832 |
| Disengagement axis | Fig. 4 | sub-456772_ses-20191122T144855 |

**Table 3.** Cross-session transfer pairs. The latent dynamics *A* and variability Σ are transferred from the source to the target session.

| Figure(s) | Source session | Target session |
| --- | --- | --- |
| Fig. S8, panel 1 | sub-456772_ses-20191120T115527 | sub-442571_ses-20190228T140832 |
| Fig. S8, panel 2,<br>Fig. 5f | sub-456772_ses-20191120T115527 | sub-484676_ses-20210416T140326 |
| Fig. S8, panel 3 | sub-480135_ses-20210227T131716 | sub-460434_ses-20191112T114755 |
| Fig. S8, panel 4 | sub-480134_ses-20210111T122809 | sub-484676_ses-20210416T140326 |

### D.2 Task inputs and coding dimensions

For the mouse ALM data, the input term *I*(*t*) in Eq. 1 encodes the task structure and movement feedback:

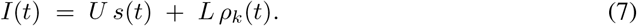

The first term is a low-dimensional task drive shared across trials, where *s*(*t*) ∈ ℝ^*P*^ consists of three brief stimulus pulses during the sample period (with separate learned vectors for left and right conditions) followed by a shared go-cue pulse. The second term injects movement-related feedback after the go cue: *ρ*_*k*_(*t*) ∈ ℝ^2^ are the measured left/right lick rates on trial *k*, and *L* ∈ ℝ^*N ×*2^ is learned. The lick inputs were intended to capture sensory and proprioceptive feedback associated with licking rather than to generate the motor output itself.

We computed coding dimensions corresponding to stimulus, choice, and response as directions in neural population activity that capture task-relevant differences between behavioral conditions [Chen et al., 2024]. Neural activity was first averaged across trials to obtain condition-specific population trajectories. A stimulus-aligned signal was defined as the difference between trial-averaged activity for the two stimulus conditions (left vs. right stimuli). To isolate choice- and response-related activity independently of stimulus, trials were grouped by behavioral choice (combining correct and error trials), and the difference between the corresponding trial-averaged trajectories (left vs. right choices) was computed.

Coding vectors were obtained by averaging these difference signals within task-relevant time windows: 0.6 s following stimulus onset (stimulus), the last 0.6 s of the delay period preceding the go cue (choice), and the first 0.6 s after the go cue (response). The resulting vectors were orthogonalized using Gram-Schmidt and normalized to unit norm, yielding three orthonormal coding dimensions in ℝ^*N*^ .

### D.2 Training

For the ALM data, our goal is to capture structured single-trial variability rather than to maximize neuron-wise fit. Full-tensor MSE on firing rates is dominated by high-variance neurons and directions of population activity, which need not be the task-relevant ones. We therefore train the inference model to match summary statistics of the activity [Pellegrino et al., 2024], including projections onto the task coding dimensions (Appendix D.2); Appendix D.4 shows that this objective better captures the stimulus and choice coding dimensions, at a modest cost in full-tensor *R*^2^.

We use two settings: an inference model, in which trial-specific biases are optimized to reconstruct recorded activity, and a generative model (Section 3.4), in which they are sampled from a learned distribution. All results use the inference setting except where noted. The inference model matches summary statistics of the single-trial population activity rather than the full activity tensor (Appendix D.4).

With 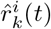 denoting model-simulated rates of neuron *i* on trial *k* at time *t*, the three data terms are:

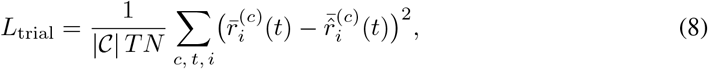

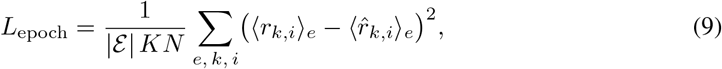

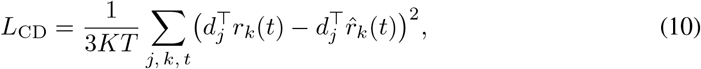

where 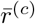 is the trial average over condition *c* ∈ {L, R}, ⟨·⟩_*e*_ denotes the average over task epoch *e* ∈ {sample, delay, response}, and 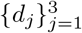 are the stimulus, choice, and response coding dimensions [Chen et al., 2024] (Appendix D.2). The full objective is

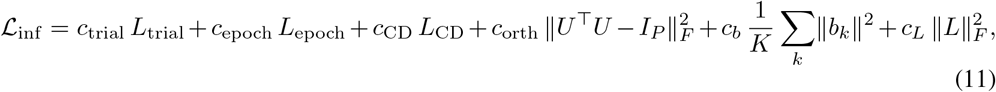

Where 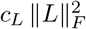 regularizes the lick input matrix. We used *c*_trial_ = 0.65, *c*_epoch_ = 0.14, *c*_CD_ = 0.02, *c*_*b*_ = 5 × 10^*−*2^, *c*_*L*_ = 0.5 and *c*_orth_ = 500, with *P* = 20, *τ* = 50 ms and Δ*t* = 10 ms.

#### Generative model

In the generative model, we draw per-trial biases from a distribution, *b*_*k*_ ∼ *N* (0, Σ), with learned Σ. Since generated trials are not aligned to recorded trials, training uses a trial-matching objective [Sourmpis et al., 2023]: for each pair within a condition, a pairwise cost *l*(*r*_*k*_, 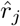) is computed as a weighted MSE over the same summary statistics as ℒ_inf_, and the optimal trial assignment is found via the Hungarian algorithm,

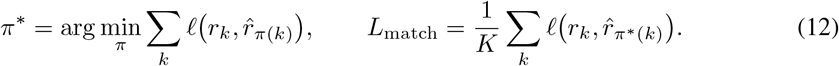

The full generative objective is

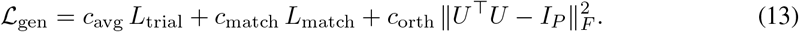

Here *c*_avg_ = 1.0, *c*_match_ = 2.0 and *c*_orth_ = 500; all other settings are as above.

### D.4 Comparison of training objectives

We compared the summary-statistics objective used in this work to direct MSE optimization on the full activity tensor. Models were trained on a single ALM session (Section 3.2) with identical architectures, hyperparameters, and train/validation splits, differing only in the training loss, and evaluated on held-out trials across multiple random initializations. The two objectives capture complementary aspects of the data: full-tensor MSE improves reconstruction of the activity tensor, while the summary-statistics objective more accurately reproduces task-aligned population structure, particularly along the stimulus and choice coding dimensions (Fig. S5).

### D.5 Reconstruction of single-trial activity

The summary-statistics objective (Appendix D.3) does not directly fit the trajectories of individual neurons on individual trials. Nevertheless, it captures single-trial responses across trial types (Fig. S6). Across all neurons, the model achieves *R*^2^ = 0.64 on the full activity tensor of held-out trials (Section 3.2).

### D.6 Disengagement axis

For each stimulus condition *c* ∈ {L, R} separately, we computed the encoding axis (Appendix B) of a binary label *y*_*k*_: 1 for ignore trials (no lick) and 0 for responded trials (correct and error). The disengagement axis is therefore the difference between the mean biases of ignore and responded trials,

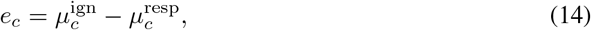

and projections onto *e*_*c*_ increase toward ignore-like trials.

Perturbations of responded trials used *y*^*′*^ = *λ*, giving

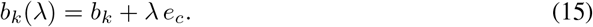

Values *λ >* 1 shift trials further along the axis than the average ignore trial. Shifts toward the responded regime were applied to ignore trials with *λ <* 0.

### D.7 Distributions of single-trial statistics in generated trials

The generative model (Section 3.4) reproduces the distribution of mean population firing rates across trials (Fig. S7a) and of single-trial projections onto the stimulus, choice, and response coding dimensions, for both left and right trials (Fig. S7b–d).

### D.8 Cross-session transfer

We compared three models on each target session: a baseline trained directly on that session; a transfer model in which the latent dynamics *A* and the bias covariance Σ were taken from a source session and held fixed while the remaining parameters were retrained; and a random control in which *A* and Σ were replaced by norm-matched random matrices. Session pairs are listed in Table 3.

## E MC Maze

### E.1 Data and preprocessing

We evaluated the model on the MC Maze dataset from the Neural Latents Benchmark (NLB) [Pei et al., 2021, Churchland et al., 2010], containing recordings from primary motor and dorsal premotor cortex of a macaque performing a delayed reaching task, spanning 108 different task configurations differing in target position and in the number and placement of barriers. In the maze reaching task [Churchland et al., 2010], a target appears in some conditions with virtual barriers; after a delay, a go cue instructs the monkey to reach to the target, producing straight or curved reaches depending on barrier placement.

Following the NLB’s standard preprocessing, trials were aligned from 250 ms before to 450 ms after movement onset and spikes were binned at 5 ms. We used the benchmark’s fixed splits: 2295 training and 574 test trials, and 137 held-in and 45 held-out neurons, where the held-out neurons were masked on test trials.

### E.2 Training and evaluation

The model receives no task input or behavioral feedback (*I*(*t*) = 0). Because the ALM objective is defined over that task’s coding dimensions, we replace the data term in Eq. 3 with a Poisson likelihood, the observation model assumed by the benchmark’s metrics. With 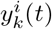 the spike count of neuron *i* on trial *k* in bin *t*,

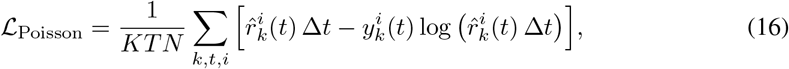

giving the full objective

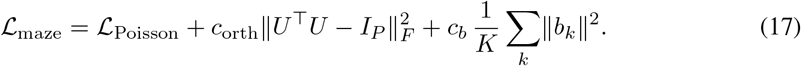

We used *c*_orth_ = 500, *c*_*b*_ = 3 × 10^*−*1^, *P* = 55 and *τ* = 40 ms. *P* and *c*_*b*_ were selected on a validation split by co-bps, over *P* from 20 to 80 and *c*_*b*_ from 10^*−*3^ to 3, with 2–3 seeds per configuration. All remaining settings are unchanged from the ALM experiments.

For evaluation we froze the trained network parameters and optimized only the per-trial biases {*b*_*k*_} of the test trials, restricting the likelihood to the held-in neurons (Appendix A). We computed all metrics with the official NLB evaluation code on the publicly released test data, and report mean ± s.e.m. over 3 random seeds.

### E.3 Neural Latents Benchmark evaluation

The benchmark results are shown in Table 4. Our model outperforms the classical baselines (smoothing, GPFA, SLDS) on all reported metrics, and falls closer to the deep sequential models (NDT, AutoLFADS), remaining below them mainly on the likelihood metrics. This is expected, as NDT and AutoLFADS are substantially more expressive. Our model is instead constrained by design: connectivity is low-rank, each recorded neuron corresponds to one network neuron, and trial-to-trial variability is restricted to a single static vector *b*_*k*_ ∈ ℝ^*P*^ within the recurrent subspace. Relative to the deep models, these constraints appear to cost single-trial, neuron-level precision while largely preserving condition-level and behaviorally relevant structure. They are also what let the inferred variability be read off as control parameters, which we analyze in Section 3.5.

**Table 4.** MC_Maze results on the held-out test set. co-bps (primary metric): likelihood of held-out neurons, in bits per spike relative to the mean firing-rates of the held-out neurons; vel *R*^2^: hand-velocity decoding from inferred rates; psth *R*^2^: match of condition-averaged rates to empirical PSTHs; fp-bps: similar to co-bps, but for held-out future timesteps of all neurons. Higher is better for all metrics. Values are mean ± s.e.m. over 3 seeds, except spike smoothing, which was run once as it is initialization-independent. Baselines from Pei et al. [2021]; dashes indicate metrics undefined for that baseline.

| Method | co-bps | vel $R^2$ | psth $R^2$ | fp-bps |
| --- | --- | --- | --- | --- |
| Smoothing | 0.211 | 0.624 | 0.183 | — |
| GPFA | $0.187 \pm 0.001$ | $0.640 \pm 0.001$ | $0.518 \pm 0.002$ | — |
| SLDS | $0.219 \pm 0.006$ | $0.775 \pm 0.006$ | $0.482 \pm 0.036$ | $-1.020 \pm 0.943$ |
| Ours | $0.283 \pm 0.002$ | $0.875 \pm 0.008$ | $0.559 \pm <0.001$ | $0.181 \pm 0.004$ |
| NDT | $0.329 \pm 0.005$ | $0.897 \pm 0.009$ | $0.579 \pm 0.023$ | $0.209 \pm 0.010$ |
| AutoLFADS | $0.346 \pm 0.005$ | $0.907 \pm 0.002$ | $0.631 \pm 0.022$ | $0.239 \pm 0.003$ |

More broadly, our aim is not optimal neuron-level reconstruction but mechanistic interpretability, something these metrics do not measure. SLDS, which scores below our model on every metric here, belongs to a family that has nonetheless supported mechanistic claims of precisely this type: rSLDS [Linderman et al., 2017] identified an approximate line attractor in mouse hypothalamus [Nair et al., 2023], and the resulting prediction about perturbing specific neurons was subsequently confirmed by targeted optogenetic stimulation [Vinograd et al., 2024].

### E.4 Reach-direction plane

In Section 3.5, we restricted the analysis to barrier-free trials, whose straight reaches are determined by the target angle *θ*_*k*_ alone (789 trials). Using the encoding regression of Appendix B, we regressed the biases on *u*(*θ*_*k*_) = (cos *θ*_*k*_, sin *θ*_*k*_)^*⊤*^:

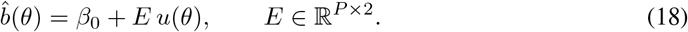

As *θ* varies, 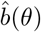 traces an ellipse in the plane spanned by the two columns of *E*, the reach-direction plane.

To change a trial’s reach direction by Δ, we moved the explained part of its bias from 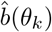 to 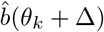, leaving the rest unchanged:

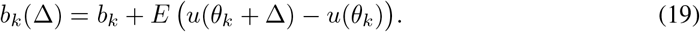

We simulated the network with the shifted biases and all other parameters fixed, and decoded hand velocity from the simulated rates with the NLB velocity decoder (ridge regression fit on the model’s rates for the training trials; Appendix E.3), kept fixed across perturbations. Each trial’s reach angle was the direction of its net hand displacement after movement onset, obtained by integrating the decoded velocity. The rotation produced by each shift was measured as the change in decoded reach angle relative to the unperturbed simulation of the same trial, averaged across trials with a circular mean.

---

**Figure S1.**
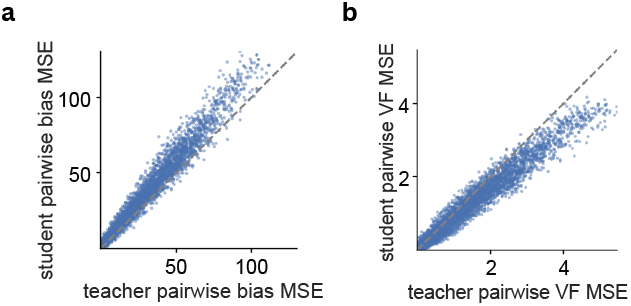
Comparison of pairwise distances between trials in the teacher and student models for the bistable system. (a) Bias space (*r* = 0.99). (b) Vector-field space (*r* = 0.99). Each point corresponds to a pair of trials, dashed lines indicate identity. The student recovers the structure of trial-to-trial variability in the teacher model.

**Figure S2.**
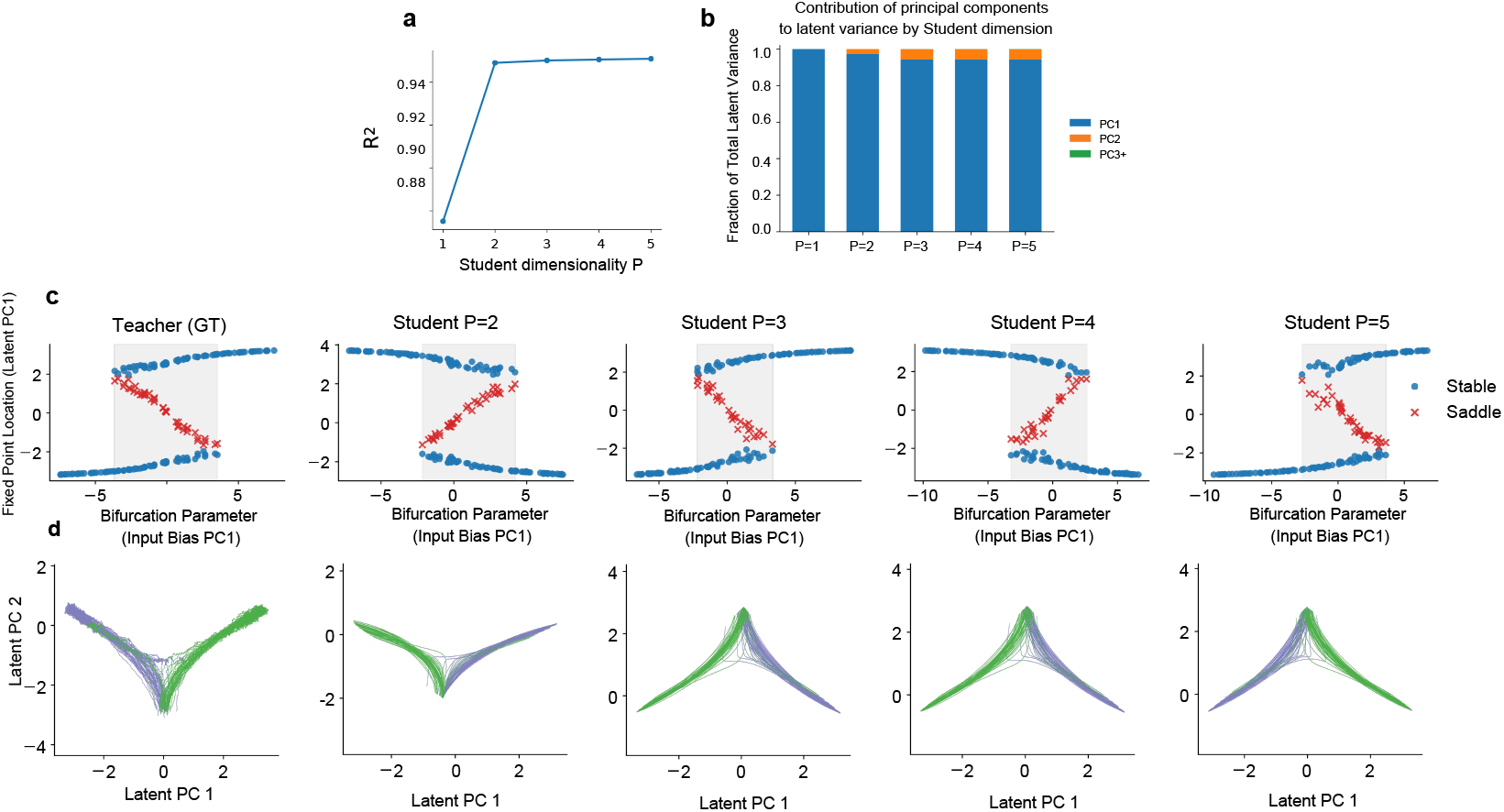
Recovery of 2D dynamics under student overparameterization (*P >* 2) for the bistable synthetic data. The student correctly identifies the underlying two-dimensional structure. (a) Firing-rate reconstruction *R*^2^ across student dimensionalities. The model’s ability to capture the observed neural rates increases and plateaus once the student reaches the true underlying dimensionality of the teacher (*P* ≥ 2) (b) Fraction of total latent variance explained by each PC. The first two PCs capture over 99% of variance for all *P*≥2, with negligible contribution from additional dimensions. (c) Recovered fixed-point locations are projected onto the first principal component of latent activity (y-axis) as a function of the first principal component of input bias (x-axis). The bifurcation diagram is preserved across all student dimensionalities. (d) Latent trajectories projected onto the top two PCs for each student. The teacher’s attractor geometry is faithfully replicated regardless of student dimensionality. Note: Because visualizations in (c) and (d) rely on internal PCA projections rather than explicit coordinate alignment, the recovered structures are subject to arbitrary rotational and sign ambiguities, though their underlying topology remains identical to the teacher.

**Figure S3.**
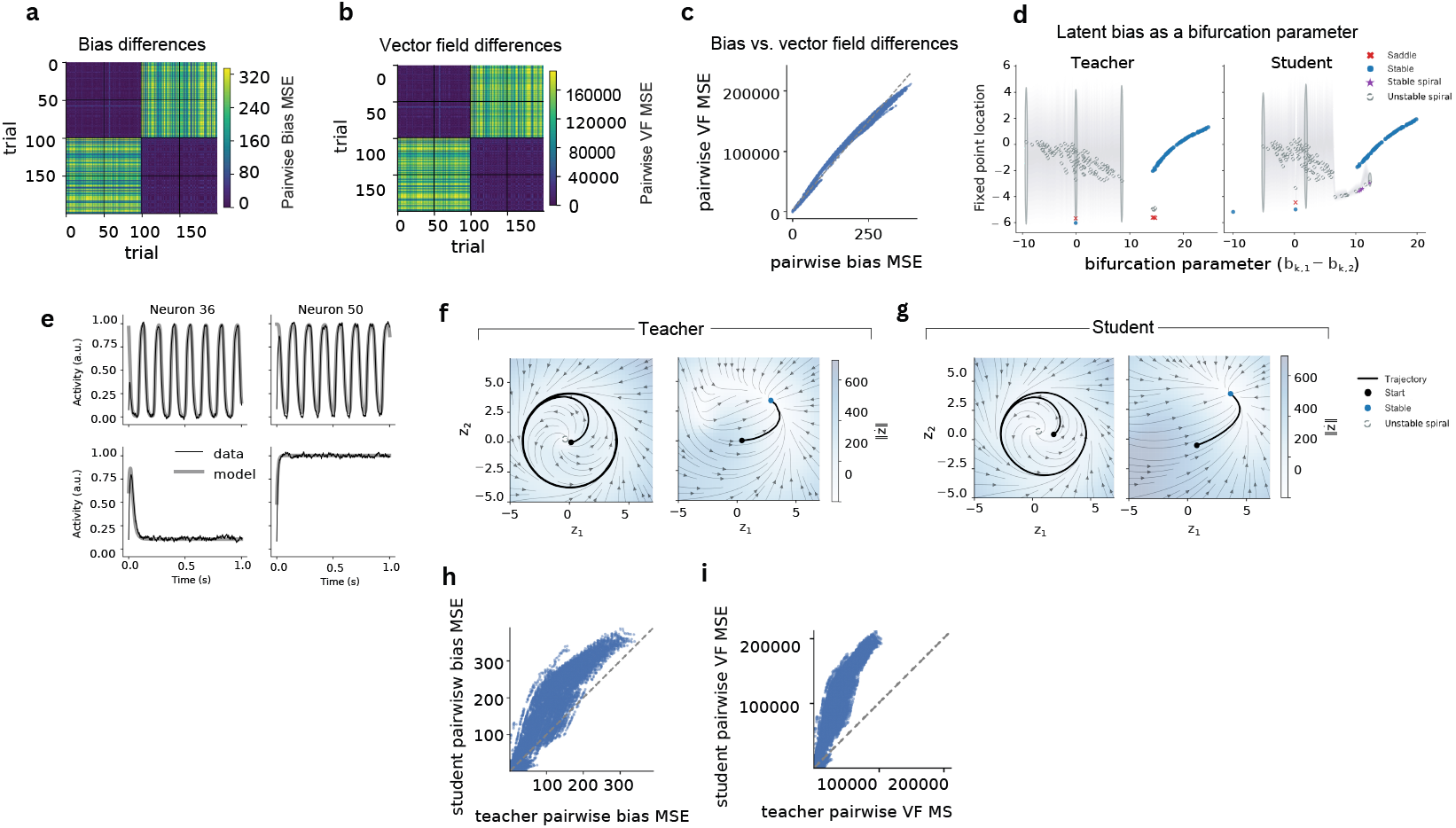
Recovery of limit-cycle dynamics in the synthetic system. (a,b) Pairwise distances between trials in bias space (a) and vector-field space (b) for the student model. (c) Pairwise differences in bias space versus the corresponding differences in vector-field space for the student model. (d) Fixed point locations as a function of bias difference. The shaded gray region shows the amplitude envelope of the limit cycle originating from unstable spirals. For visualization, the latent spaces are aligned using a Procrustes transformation. (e) Single-neuron activity for two representative trials: oscillatory (left) and fixed-point (right). (f,g) Vector fields for representative trials for the teacher (f) and student (g) models, after Procrustes alignment (h,i) Pairwise trial distances in bias space (h) and vector-field space (i) for teacher vs. student (*r* = 0.95; dashed line: identity).

**Figure S4.**
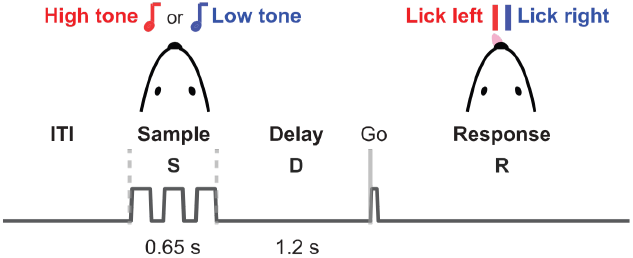
Delayed-response task in mouse ALM (reproduced from Chen et al. [2024]). One of two auditory tones presented during the sample epoch (S; 0.65 s) instructs a left or right lick. After a delay (D; 1.2 s), a go cue marks the start of the response epoch (R), in which the mouse reports its choice by licking.

**Figure S5.**
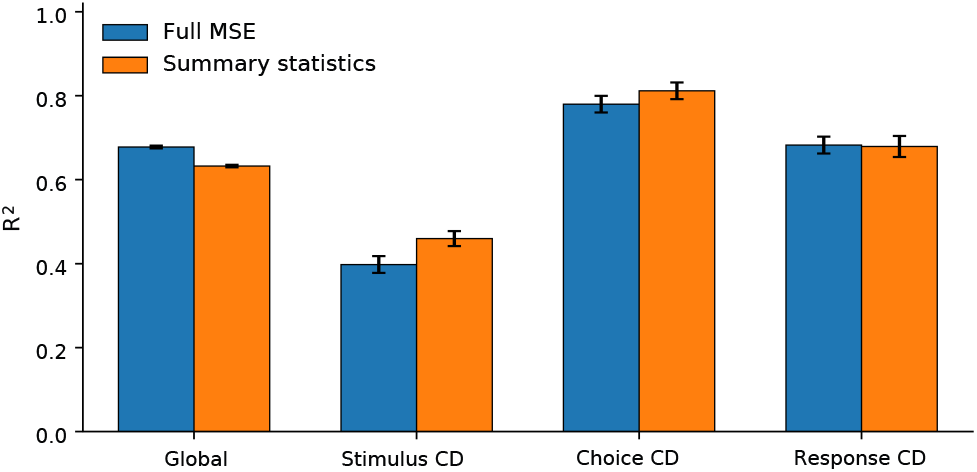
Comparison of model performance under two training objectives. Bars show mean *R*^2^ on held-out validation trials for models trained with full-tensor MSE (blue) and the summary-statistics objective of Appendix D.3 (orange), averaged over random seeds; error bars indicate standard deviation across seeds. Metrics include *R*^2^ on the full activity tensor and *R*^2^ of projections onto the stimulus, choice, and response coding dimensions.

**Figure S6.**
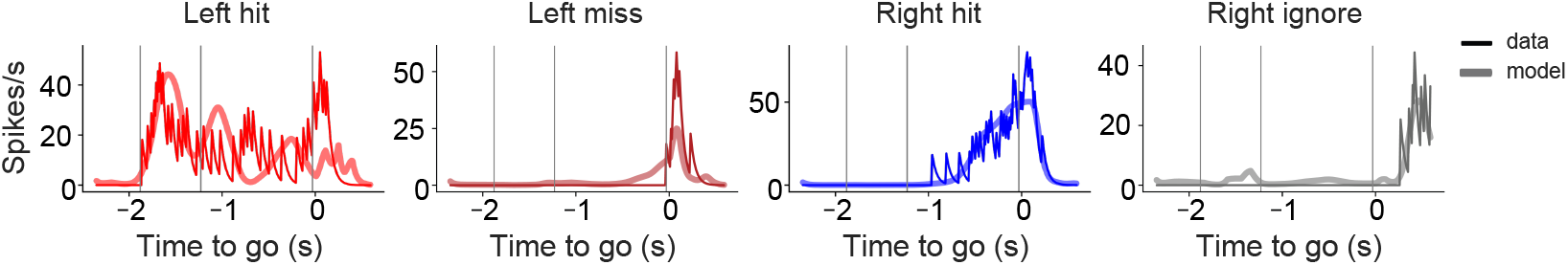
Single-trial activity of example ALM neurons across trial types (dark: data; light: model). Session as in Fig. 3.

**Figure S7.**
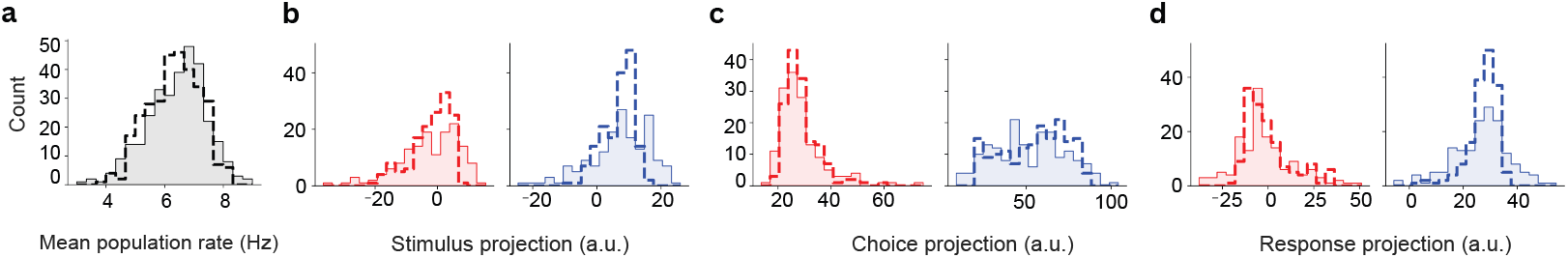
Distributions of single-trial statistics for held-out recorded trials (solid) and generated trials (dashed). (a) Mean population firing rate, averaged across neurons and time within each trial. (b–d) Single-trial projections onto the stimulus (b), choice (c), and response (d) coding dimensions, averaged over the corresponding task epoch, for left (red) and right (blue) trials. Session as in Fig. 5c,d.

**Figure S8.**
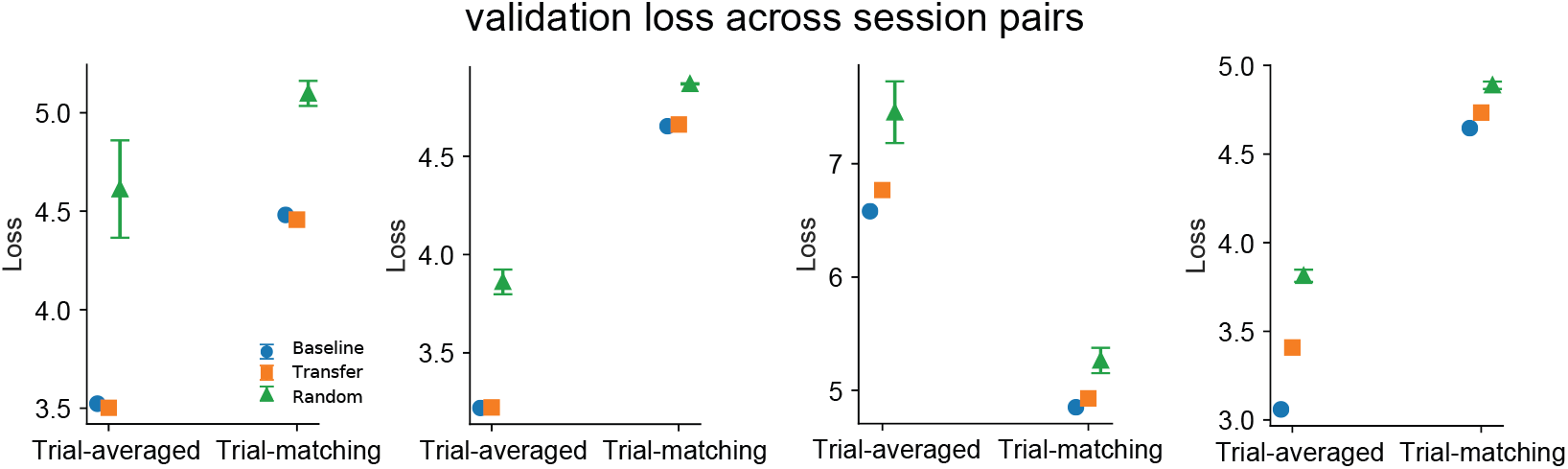
Cross-session transfer performance across multiple session pairs. Validation losses for baseline, transfer, and random control models are shown for both trial-averaged and trial-matching objectives. Across session pairs, transfer models consistently achieved performance close to baseline models and substantially better than random controls, indicating that the learned latent dynamics and variability structure generalize across sessions.

